# Substrate Specificities of DDX1: A Human DEAD-box protein

**DOI:** 10.1101/2024.01.09.573566

**Authors:** Anthony F. T. Moore, Yepeth Berhie, Isaac S. Weislow, Eda Koculi

**Affiliations:** Department of Chemistry, University of Central Florida, 4111 Libra Drive, Physical Sciences, Orlando, FL 32816-2366; Department of Chemistry and Biochemistry, The University of Texas at El Paso, 500 W University Ave, Chemistry and Computer Science, El Paso, TX, 79902-5802

## Abstract

DDX1 is a human protein which belongs to the DEAD-box protein family of enzymes and is involved in various stages of RNA metabolism from transcription to decay. Many members of the DEAD-box family of enzymes use the energy of ATP binding and hydrolysis to perform their cellular functions. On the other hand, a few members of the DEAD-box family of enzymes bind and/or hydrolyze other nucleotides in addition to ATP. Furthermore, the ATPase activity of DEAD-box family members is stimulated differently by nucleic acids of various structures. The identity of the nucleotides that the DDX1 hydrolyzes and the structure of the nucleic acids upon which it acts in the cell remain largely unknown. Identifying the DDX1 protein’s *in vitro* substrates is important for deciphering the molecular roles of DDX1 in cells. Here we identify the nucleic acid sequences and structures supporting the nucleotide hydrolysis activity of DDX1 and its nucleotide specificity. Our data demonstrate that the DDX1 protein hydrolyzes only ATP and deoxy-ATP in the presence of RNA. The ATP hydrolysis activity of DDX1 is stimulated by multiple molecules: single-stranded RNA molecules as short as ten nucleotides, a blunt-ended double-stranded RNA molecule, a hybrid of a double-stranded DNA-RNA molecule, and a single-stranded DNA molecule. Under our experimental conditions, the single-stranded DNA molecule stimulates the ATPase activity of DDX1 at a significantly reduced extent when compared to the other investigated RNA constructs or the hybrid double-stranded DNA/RNA molecule.

## INTRODUCTION

Based on their conserved amino acid motifs, helicases are segregated in five super families^1^. DEAD-box proteins, named after their conserved amino acid sequences, make up a large number of Super Family 2 members ^1^. The human DDX1 protein is a member of the DEAD-box enzyme family, which engages in a wide range of cellular processes, including DNA double-strand break repair, RNA transcription, ribosomal RNA (rRNA), microRNA (miRNA) and transfer RNA (tRNA) processing, messenger RNA (mRNA) 3’ end maturation, translation initiation, and RNA transport, storage, and decay^2–13^. Consequently, this protein is implicated in the progressions of cancers, viral infections, and embryonic development^3, 14–19^. Here we examine the identity of the nucleotides that the DDX1 protein hydrolyzes and the properties of the structures and sequences of the nucleic acid substrates that support the DDX1 protein’s ATPase activity. This knowledge could inform how the DDX1 protein performs its cellular functions.

All DEAD-box proteins possess two covalently linked RecA-like domains which form the nucleotides, nucleic acid binding pockets and regions of communication between these pockets^20–23^. Many members of the DEAD-box RNA family use the energy of ATP and hydrolysis to unwind short RNA double helixes, perform clamping, and display proteins from RNA and/or anneal RNA^22–24^. With a few exceptions, the ATP hydrolysis activity of DEAD-box proteins is stimulated by RNA molecules^22, 25^. DEAD-box proteins are RNA-dependent ATPases^22, 25^. The exceptions are the human DEAD-box proteins DDX3X^26, 27^, DDX43^28^, Dbp5p (DDX19)^29^, *Saccharomyces cerevisiae* (*S. cerevisiae*) DEAD-box proteins Dbp5p^29^ and Dbp9p^30^, and the *Pisum sativum* DEAD-box protein p68^31^. ATP hydrolysis activities of the latter DEAD-box proteins are stimulated by both RNA and DNA.

Exceptions are also noted on the nucleotide substrate preferences of the DEAD-box RNA proteins. Although most of the investigated DEAD-box proteins hydrolyze only ATP and deoxy ATP^22, 25, 32^, the human DEAD-box protein DDX3X^26, 33^ and the *S. cerevisiae* DEAD*-*box protein Mss166^34^ bind and hydrolyze all four nucleotides and deoxynucleotides. In addition, the *S. cerevisiae* DEAD-box protein Dhh1 binds all four nucleotides but is unable to hydrolyze these nucleotides^35^.

Here, taking advantage of Malachite Green^36, 37^ and Thin Layer Chromatography (TLC)^37, 38^ assays, we systematically examined whether the DDX1 protein hydrolyzes additional nucleotides in addition to ATP and deoxy-ATP. Moreover, we examined the RNA sequence and structural properties required to support the DDX1 protein ATP hydrolysis activity. Lastly, we investigated whether a single-stranded DNA (ssDNA) molecule and a blunt-ended RNA/DNA hybrid double helix (dsRNA/DNA) support the ATP hydrolysis activity of DDX1.

The human DDX1 protein is 720 amino acids long^5, 18, 39^. The human DDX1 protein construct employed in this study lacks 12 amino acids from its C-terminal end. The 12 C-terminal residues are predicted to be unstructured and outside the RecA-like C-terminal domain region^39^. The DDX1 construct lacking the 12 terminal amino acids residues has been used successfully in previous studies to investigate the DDX1 protein’s catalytic cycle and the ability of the DDX1 protein to convert G quadruplexes to R-loops structures ^8, 39^.

## MATERIALS AND METHODS

### Materials

2-Mercaptoethanol (BME), Bradford reagents, and glacial acetic acid were obtained from Sigma-Aldrich. DTT, Tris-base, concentrated HCl (12.1 M), MgCl_2_, KCl, KOH, 40% acrylamide:bis-acrylamide solution (29:1 and 37.5:1), glycerol, and urea were obtained from Fisher. Phenylmethylsulfonyl fluoride (PMSF) was obtained from Biosynth. The random RNA and/or DNA strands, as well as the poly(A)_10_, poly(C)_10_, poly(G)_10_ and poly(U)_10_ RNAs used in ATPase activity analysis, were purchased from Integrated DNA Technologies, Inc. (Coralville, IA, USA). [γ-^32^P]-ATP (6000 Ci/mmol, 150 mCi/mL) was purchased from Perkin Elmer. T4 polynucleotide kinase, CTP, GTP, UTP, dATP, dCTP, dGTP, and dTTP were purchased from New England Biolabs. The tRNA^Phe^ from *S. cerevisiae* was purchased from Sigma-Aldrich.

### DDX1 Expression and purification

The DDX1 clone was a gift from the Roseline Godbout laboratory at the University of Alberta^5^. The coding sequence of the DDX1 protein was PCR amplified and inserted into the pET28 vector (Novagen). A His-tag was placed in the N-terminal of DDX1 and a stop codon was inserted at the coding sequence of amino acid 708. The insertion of the stop codon created a 12 amino acid truncated DDX1 construct. The truncated construct expressed at a much higher yield in *E. coli* cells than in the intact protein (data not shown).

*E. coli* BL21 (DE3) CodonPlus was transformed with the pET28a vector bearing the coding sequence of the DDX1 protein, and cells were grown in Luria Broth media containing 30 μg/mL of Kanamycin and 34 μg/mL Chloramphenicol. Purification of the expressed DDX1 was carried out similar to procedures published by Kellner, *et al*.^40^ Briefly, cell pellets were resuspended in a nickel column equilibrating buffer (50 mM Tris-HCl pH 8.0 at 4 °C, 250 mM KCl, 10 mM BME, and 1 mM PMSF) and supplemented with a protease inhibitor cocktail (cOmplete Protease Inhibitor Cocktail Tablets, EASYpack, Roche). Cells were lysed by sonication on ice (Branson Digital Sonifier 450, equipped with a standard flat tip, 0.5 in outer diameter, set at 50% amplitude) in six 30s pulses, 5 min between pulses. The lysate was spun down at 14,000 rpm for 60 min. The supernatant was loaded onto an 8 mL Ni-NTA column. The protein was eluted with 250 mM imidazole. Pooled fractions were diluted 1:10 with a heparin equilibrating buffer (50 mM Tris-HCl pH 8.0 at 4 °C, 5 mM MgCl_2_, 3 mM DTT, and 1 mM PMSF), loaded onto a 5 mL Heparin column (HiTrap Heparin HP, GE Healthcare Life Sciences), and washed with a heparin dilution buffer supplemented with 10 mM KCl. The protein was eluted on a linear gradient with a heparin dilution buffer supplemented with 1 M KCl. Fractions containing the desired protein were pooled and diluted 1:10 with an anion-exchange dilution buffer (50 mM Tris-HCl pH 9.0 at 4 °C, 5 mM MgCl_2_, 3 mM DTT, and 1 mM PMSF), loaded onto an anion-exchange column (HiTrap Q HP, GE Healthcare Life Sciences), washed with an anion-exchange dilution buffer supplemented with 30 mM KCl, and eluted with a linear gradient of an anion-exchange dilution buffer supplemented with 1 M KCl. Fractions containing the desired protein were pooled, concentrated, and further purified via a gel-filtration column (Superdex 200 10/300 GL, GE Healthcare), which was equilibrated with an RNase-free gel-filtration buffer (10 mM HEPES-KOH pH 8.0, 250 mM KCl, 5 mM MgCl_2_, 3 mM DTT). Protein concentrations were determined via a Bradford assay using SpectraMax Plus 384 (Molecular Devices). Pooled fractions were concentrated via 10 kDa centrifugal filters (Amicon Ultra Ultracel-10K, Millipore) and stored at −80°C in small aliquots.

### Malachite Green Phosphate assay

The Malachite Green-molybdate assay was employed to measure the DDX1 protein’s NTP and deoxy-NTP nucleotide hydrolysis activity ^36, 37^. The Malachite Green–molybdate reagent was prepared by the mixing 0.034% of malachite green, 1.04% of ammonium molybdate, and 1 M HCl stirring constantly overnight at 4°C, filtering the solution, and storing it at 4°C. Polyvinyl alcohol, which acts as a stabilizing reagent, was added to a concentration of 0.4%. The final mixture was equilibrated at room temperature before use.

The nucleotide and deoxynucleotide hydrolysis reactions were performed in a buffer consisting of 50 mM Tris-HCl (pH 8.0), 250 mM KCl, 2 mM MgCl_2_, 2 mM DTT, 20 μM tRNA^Phe^, 1 μM DDX1, and 1 mM nucleotide or deoxynucleotide. The control reactions lacked nucleotides, deoxynucleotides, DDX1 or tRNA^Phe^. Reactions were initiated by the addition of a nucleotide or a deoxynucleotide solutions. 10 μL of DDX1 hydrolysis reactions were allowed to continue for 60 min at 23 °C in 384-well microplates. The hydrolysis reaction was quenched with 2.5 μL of 50 mM EDTA-KOH (pH 8.0) and incubated for an additional 5 minutes before adding 75 μL of the Malachite Green-molybdate reagent. After 1 min, the Malachite Green-molybdate reagent reaction was quenched with 10 μL of 34% sodium citrate and allowed to develop for 50 min, before absorbance was measured at 625 nm using SpectraMax Plus 384 (Molecular Devices).

### TLC ATPase assay

Hydrolysis of ATP was monitored using TLC as previously described ^37, 38, 41^. First, the RNA was annealed, if necessary, by incubating it with 50 mM HEPES-KOH pH 7.5, 50 mM KCl at 95 °C for 1 min, 65 °C for 3 min, and cooled down to 25 °C for 1 min, before adding MgCl_2_ to a final concentration of 10 mM. The annealed solution was incubated at 23 °C for 20 min before placing on ice. Subsequently, varying concentrations of the RNA substrate were mixed with 25 mM Tris-HCl pH 8.0, 2 mM MgCl_2_, 250 mM KCl, 2 mM DTT, 0.5 mM EDTA-KOH pH 8.0, 1 μM DDX1, 6 mM ATP, and 2.5 nM γ-^32^P-labeled ATP, then incubated at 37 °C for 60 min. After incubation, the reactions were quenched with 0.5 M EDTA pH 8 and spotted on PEI-Cellulose F coated TLC plates. The TLC plates were developed using a solvent system consisting of 0.75 M LiCl and 1 M acetic acid. TLC plates were left to dry, then exposed to phosphor imaging screens. The screens were scanned using a Personal Molecular Imager System (Bio-Rad Laboratories) and analyzed using Quantity One (Bio-Rad Laboratories). The fraction of ATP hydrolyzed was calculated as the ratio of the counts on the inorganic phosphate band over the total counts on the lane. The fraction of ATP hydrolyzed versus protein concentrations were fit to the Michaelis-Menten equation (Equation 1) using OriginLab:

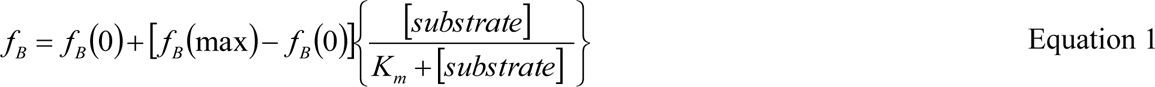

where *f_B_* is the maximum amount of ATP hydrolyzed, *f_B_(0)* and *f_B_(max)* are the lower and upper baselines of the ATP hydrolysis, *K_m_* is the Michaelis constant and [substrate] is the nucleic acid concentration.

## RESULTS AND DISCUSSION

### DDX1 hydrolyzes ATP and deoxy-ATP only in the presence of tRNA^Phe^, but it is unable to hydrolyze other nucleosides

The specificity of the DDX1 protein for different nucleotides and deoxynucleotides was examined by employing the Malachite Green Phosphate assay^36, 37^. This assay quantifies the complex formed between Malachite Green, molybdate, and the inorganic phosphate produced by the DDX1 protein hydrolysis of nucleotides or deoxynucleotides. The assay was performed in the presence or absence of the DDX1 protein, tRNA^Phe^, nucleotide, and deoxynucleotide (Figure 1). Our data show that the DDX1 protein, in the presence of tRNA^Phe^, hydrolyzes only ATP and deoxy-ATP. The DDX1 protein is unable to hydrolyze other nucleotides or deoxynucleotides.

**Figure 1.**
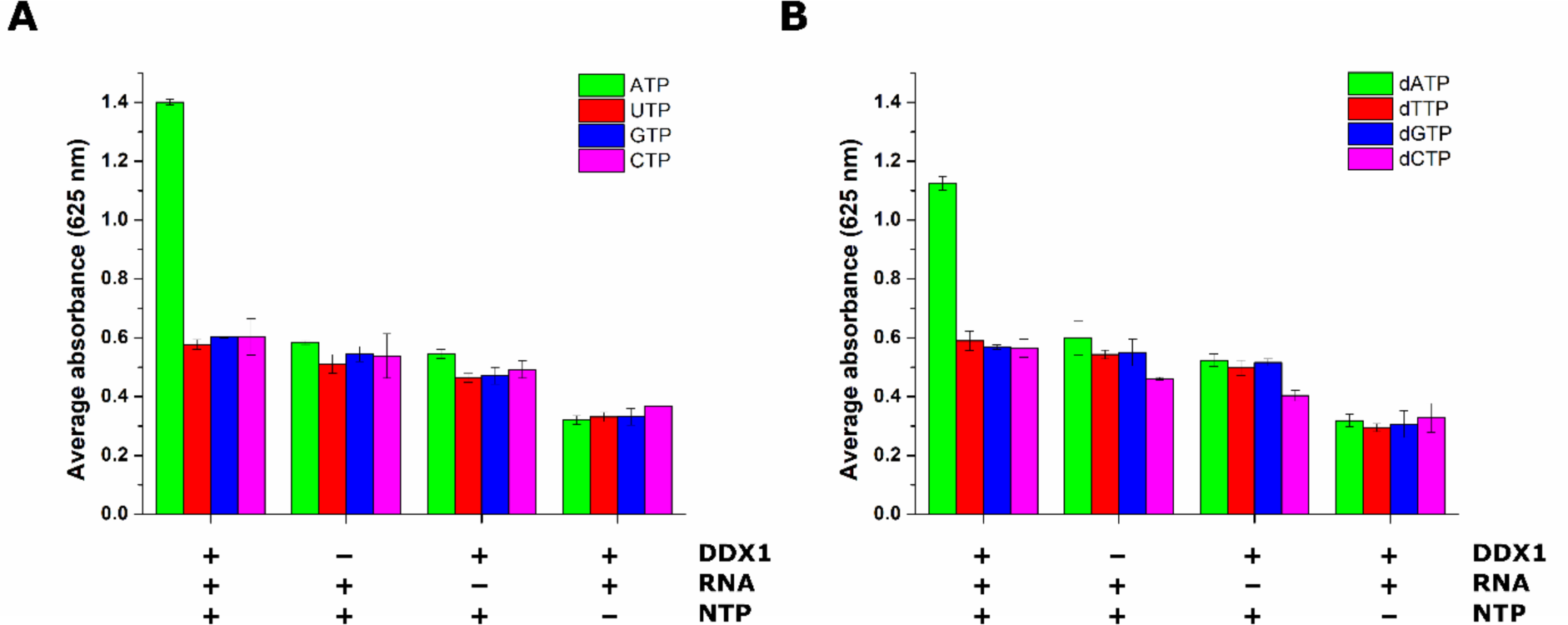
The DDX1 protein hydrolyzes ATP and dATP in the presence of tRNA^Phe^ from *S. cerevisiae.* Malachite Green colorimetric assay was employed to examine the DDX1 protein specificity for the four nucleotides and deoxynucleotides. (A) Nucleotides hydrolyzed by DDX1 in presence of tRNA: ATP (green), UTP (red), GTP (blue), and CTP (purple); and (B) Deoxynucleotides hydrolyzed by DDX1 in the presence of tRNA^Phe^: dATP (green), dTTP (red), dGTP (blue), and dCTP (purple). The values and error bars represent the averages and the standard deviations computed from three independent experiments.

### A 10-nucleotide long RNA substrate is sufficient for supporting the ATPase activity of DDX1

An investigation of different DEAD-box protein crystal structures in complex with single-stranded RNA reveals that the footprint of the DEAD-box protein catalytic core on RNA is between 6-10 nucleotides^42–45^. Moreover, the Dhh1 protein binds a 12-nucleotide long poly(U) 15-fold tighter than a 10-nucleotide long poly(U), while it binds a 20-nucleotide long poly(U) 30-fold tighter than a 10-nucleotide long poly(U)^35^. Thus, DEAD-box proteins require different single-stranded RNA lengths for tight complex formation and/or stimulation of the ATPase activity. Here we investigate the single-stranded RNA length requirement for supporting the DDX1 protein ATP hydrolysis activity.

The TLC assay was employed to determine the RNA length required to support the DDX1 protein ATP hydrolysis activity. TLC has been used extensively to examine the stimulation of the DEAD-box proteins’ ATP hydrolysis activity by various nucleic acid substrates^37, 38, 41^. For the experiments outlined here, we employed single-stranded RNA (ssRNA) molecules of similar sequences and different lengths. The lengths of the molecules were 10-nucleotide long, 18-nucleotide long and 36-nucleotide long (Table 1). The first ten residues in the 18-nucleotide long RNA molecule are the same as in the 10-nucleotide long RNA molecule, while the 36-nucleotide long RNA construct is the repeat of the 18-nucleotide long RNA molecule. Lastly, the Mfold was used to investigate the propensity of the 10-nucleotide long, 18-nucelotide long and 36 nucleotide long molecules to form secondary structures^46^. Mfold predicted no stable secondary structure formation for the above RNA molecules. Thus, the 10-nucleotide long, 18-nucleotide long and 36-nucleotide long molecules employed here are likely single stranded in solution.

**Table 1.**
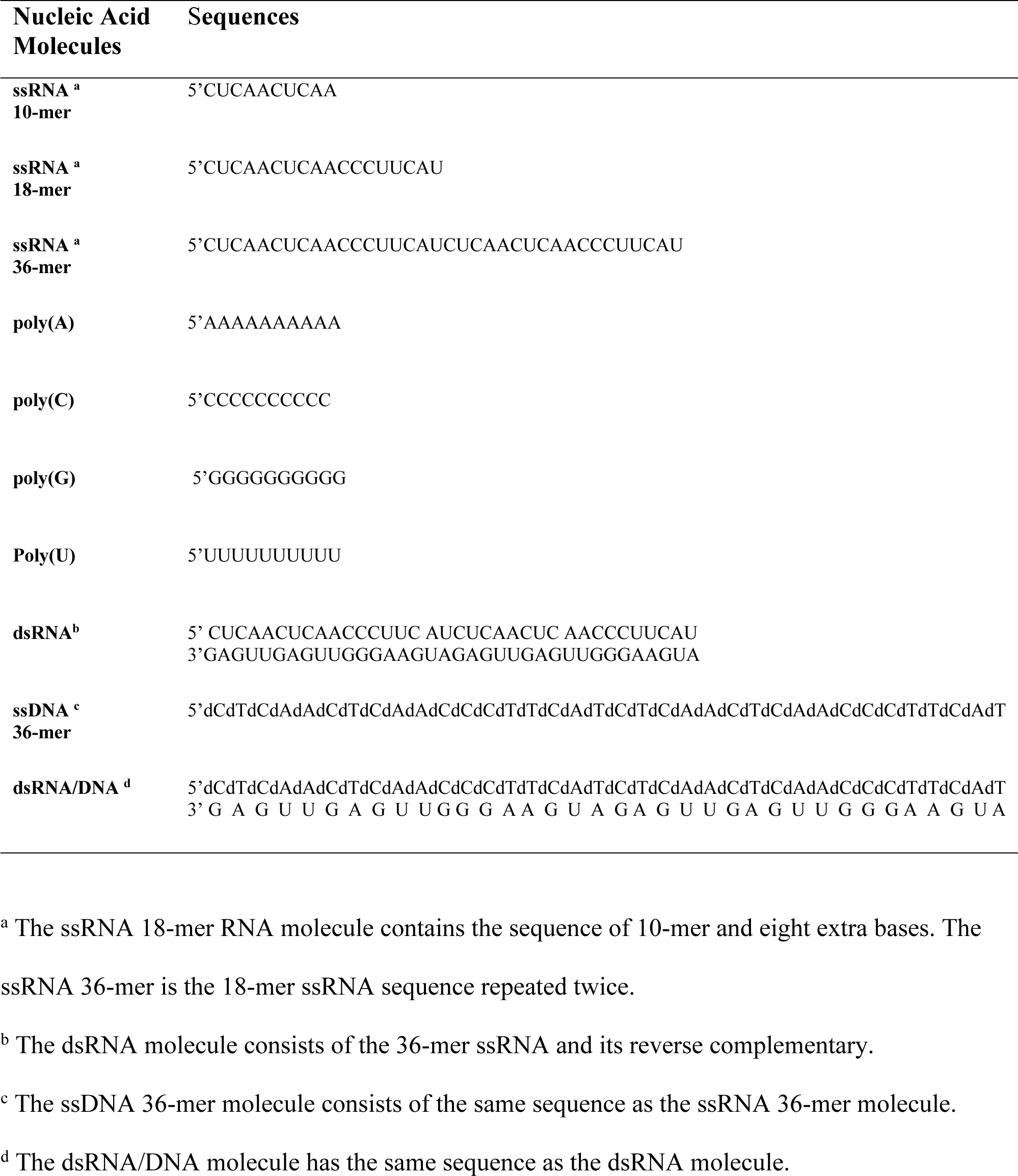
Nucleic Acid Molecules Employed for the TLC ATPase assay.

The dependence of ATP hydrolysis on the concentration of RNA, as measured by TLC, reveals that all three RNA constructs, independent of their lengths, stimulate the ATPase activity of DDX1 at the same extent and with a similar K_M_ (Figure 2, Table 2). Thus, a 10-nucleotide long RNA construct is sufficient to support the ATPase activity of DDX1.

**Figure 2.**
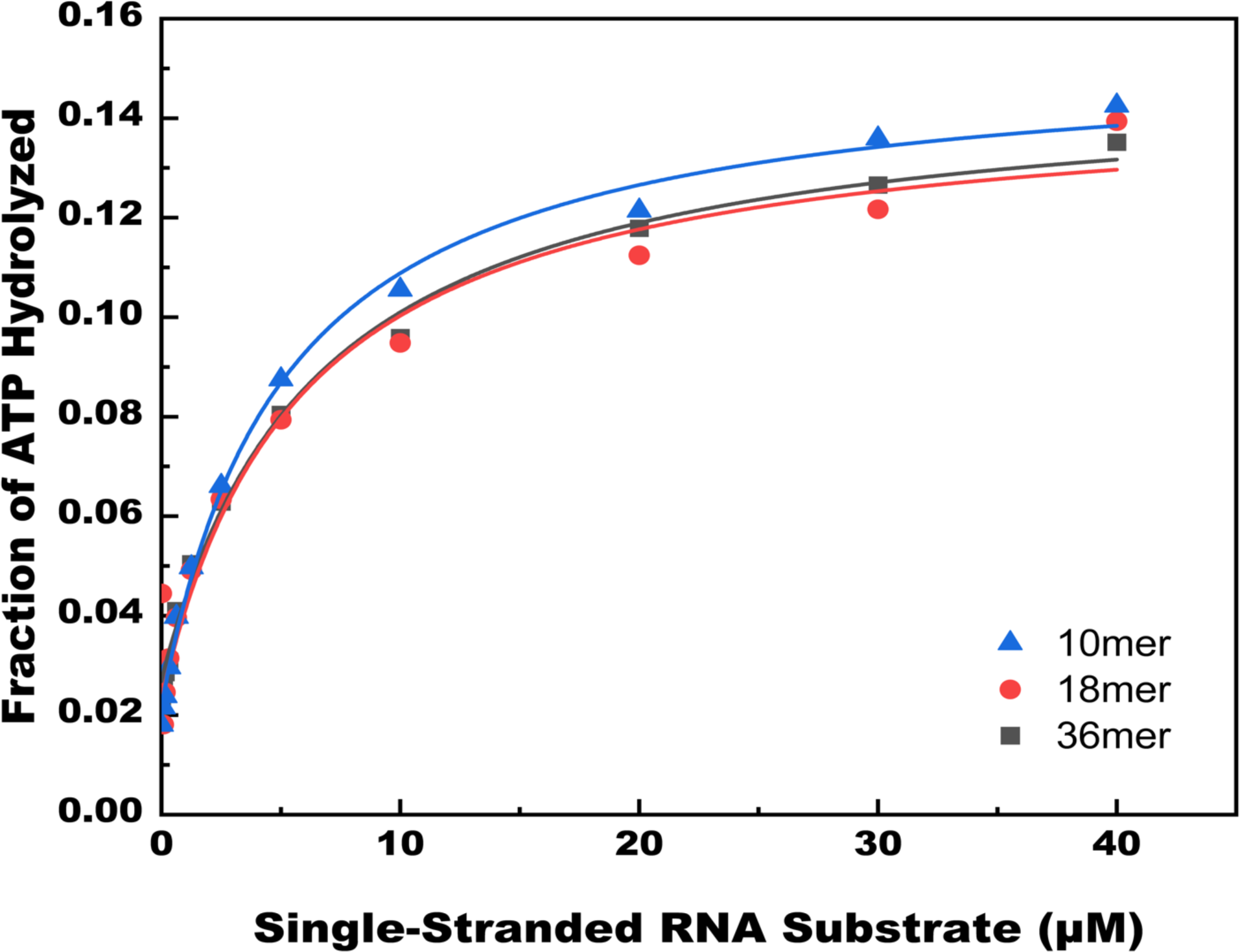
The single-stranded 10-nucleotide long RNA substrate supports the ATPase activity of the DDX1 protein to the same extent as the longer single-stranded RNA substrates. TLC was employed to determine the dependence of the DDX1 protein’s ATP hydrolysis activity on RNA concentration. The graphs represent the dependence of the DDX1 protein ATPase activity on RNA concentration. Equation 1 was used to fit the data. The average value for K_m_ and the extent of maximum ATP hydrolyzed obtained from at least two independent experiments are shown in Table 2. The sequence of the single-stranded 10-, 18- and 38-nucleotide long RNA molecules are shown in Table 1. The dependence of DDX1 ATP hydrolysis versus the 10-nucleotide long RNA concentration is shown in blue, the 18-nucleotide long RNA in red, and the 36-nucleotide long in gray.

**Table 2.** Stimulation of ATP Hydrolysis by RNA and DNA Constructs.

| **Nucleic Acid Molecules** | **K<sub>m</sub> (μM) <sup>a</sup>** | **Maximum Fraction <sup>b</sup> of ATP Hydrolyzed** |
| --- | --- | --- |
| **ssRNA 10-mer** | 5.14 ± 0.10 | 0.12 ± 0.01 |
| **ssRNA 18-mer** | 8.15 ± 4.78 | 0.15 ± 0.027 |
| **ssRNA 36-mer** | 6.10 ± 0.46 | 0.12 ± 0.003 |
| **poly(A)** | 6.06 ± 0.56 | 0.11 ± 0.004 |
| **poly(C)** | 7.68 ± 0.71 | 0.12 ± 0.004 |
| **poly(G)<sup>c</sup>** | ————— | ————— |
| **Poly(U)** | 15.58 ± 6.33 | 0.11 ± 0.004 |
| **dsRNA** | 0.92 ± 0.62 | 0.18 ± 0.032 |
| **ssDNA 36-mer** | 24.92 ± 11.95 | 0.054 ± 0.001 |
| **dsRNA/DNA** | 0.55 ± 0.06 | 0.065 ± 0.004 |

<sup>a</sup> The ATP hydrolysis in the presence of different nucleic acid molecules was determined using TLC. The K<sub>m</sub> of ATP hydrolysis was calculated using Equation 1 in the Materials and Methods section of the paper. The data shown are the averages obtained from at least two independent experiments. The error bars are the standard deviations from the averages.
<sup>b</sup> The maximum fraction of ATP hydrolyzed was obtained by fitting the Equation 1 in Materials and Methods to TLC data. The data shown are the averages and the standard deviations obtained from at least two independent experiments.
<sup>c</sup> The poly(G) molecule does not stimulate the ATPase activity of the DDX1 protein.

### 10-nucleotide long poly(A), poly(C) and poly(U) molecules support the ATPase activity of the DDX1 protein, while a 10-nucleotide long poly(G) molecule does not

Most DEAD-box proteins show sequence non-specific RNA dependent ATPase activity^22, 25, 47–50^. However, the *Thermus thermophilus* (*T. thermophilus)* DEAD-box protein Hera is unable to form the closed ATPase-competent conformation in presence of poly(G)^51^. Moreover, poly(G) and poly(U) molecules stimulate the ATPase activity of the *S. cerevisiae* DEAD-box protein Ded1p at a significantly reduced extent than the poly(A) and poly(C) molecules^52^. Lastly, poly(A), poly(C) and poly(U) stimulate the ATPase activity of Dhh1, *S. cerevisiae* Dbp5, and human Dbp5 (DDX19) at higher extent than poly(G) molecules^29, 35^. Here we investigate the DDX1 protein preference for different RNA homopolymers.

TLC was employed to investigate the preference of DDX1 for 10-nucleotide long RNA homopolymers (Figure 3, Table 1, Table 2,). All the RNA homopolymers with the exception of poly(G) support the ATPase activity of DDX1. Similarly, Kellner *et al*. found that both 10-nucleotide long poly(A) and poly(U) support the ATPase activity of DDX1^39^. 10-nucleotide long poly(C) and poly(G) constructs were not investigated in the Kellner *et al*. study^39^.

**Figure 3.**
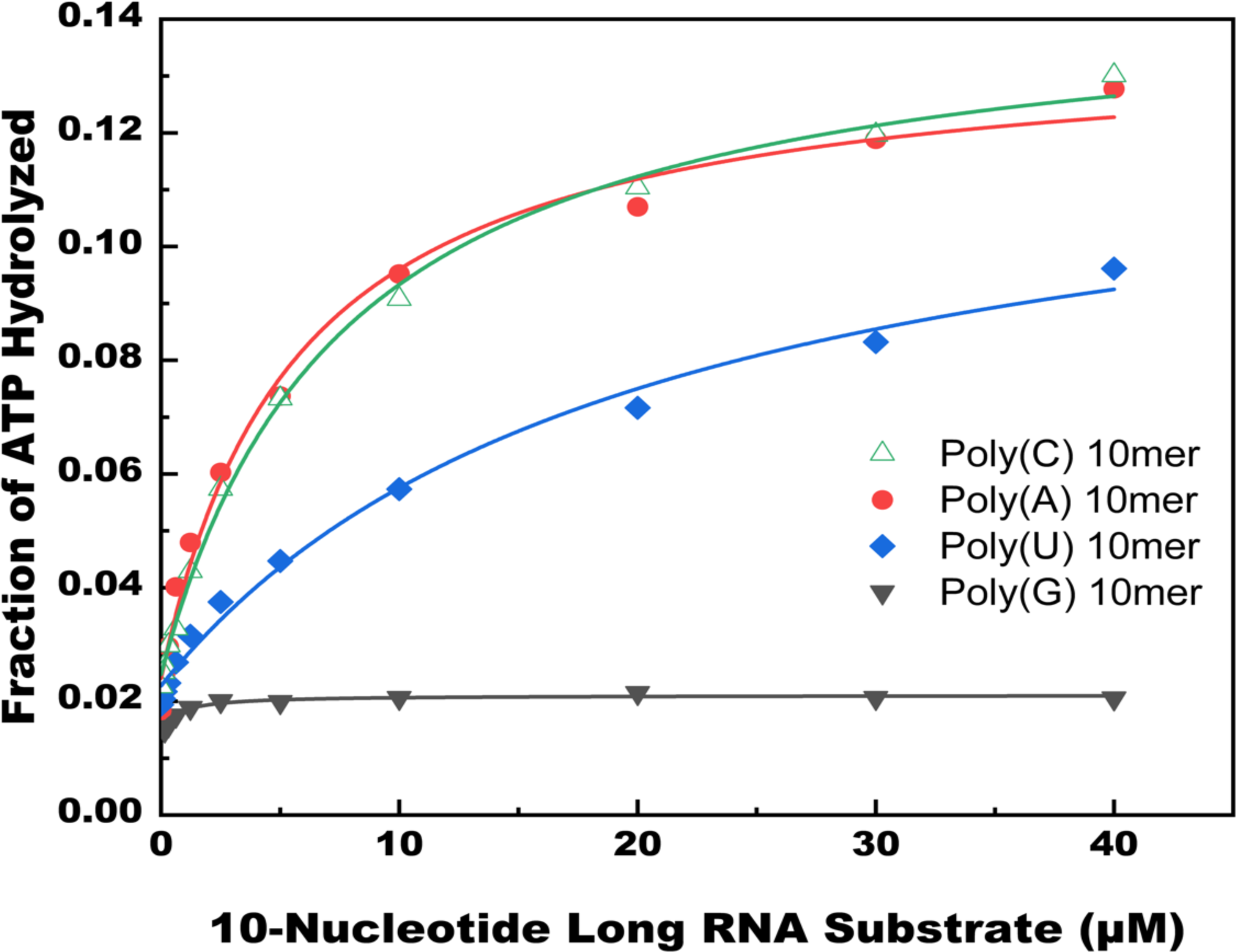
The dependence of the DDX1 protein ATPase activity on the RNA homo-oligonucleotides. TLC was employed to determine the dependence of the DDX1 ATP hydrolysis activity on 10 nucleotide-long poly(A), poly(C), poly(G) and poly(U) concentrations. The graphs represent the dependence of the DDX1 protein ATPase activity on 10 nucleotide-long poly(A), poly(C), poly(G) and poly(U) concentrations. Equation 1 was used to fit the data and determine the K_m_ and the maximum fraction of ATP hydrolyzed. The average values for K_m_ and the maximum fraction of ATP hydrolyzed obtained from at least two independent experiments are shown in Table 2. The dependence of DDX1 ATP hydrolysis versus poly(A) RNA concentration is shown in red circles, poly(C) in green triangles, poly(G) in gray triangles, poly(U) blue diamonds.

Chen *at al*. found that poly(A), poly(C), poly(U) of different lengths, from hundreds to thousands, support the ATPase activity of DDX1, while poly(G) molecules of several hundreds to thousands nucleotide lengths do not support the ATPase activity of DDX1^2^. Long poly(G) molecules form G-quadruplexes in the presence of Na^+^, K^+^ or Li^+^ ions^53^. However, Chen *at al*. did not use Na^+^, K^+^ or Li^+^ ions in their ATP hydrolysis assay, thus the poly(G) molecules in the above study could not form G-quadruplexes. QGRS Mapper predicts, due to its small size, the 10-nucleotide long poly(G) molecule employed in our study is also unable to form a G-quadruplex^54^. Therefore, both the poly(G) RNA molecules from Chen *at al*. ^2^ and the poly(G) RNA molecule from this study, which are unable to form a G-quadruplex, do not support the ATP hydrolysis activity of DDX1.

Interestingly, the DDX1 protein was shown to convert G-quadruplexes to R-loops both *in vivo* and *in vitro*, in an ATP-dependent manner^8^. Moreover, the DDX1 protein *in vitro* bound to a G-quadruplex but not to an RNA sequence rich in G nucleotides, which was unable to form the G-quadruplex structure^8^. An attractive hypothesis, based on our data and previous results, is that DDX1 has evolved to recognize G-quadruplex structures and interacts poorly with G-rich sequences unable to form G-quadruplexes.

The ability to discriminate between different nucleic acid conformations has been demonstrated for another DEAD-box protein, Mss116^34^. The A form RNA and DNA double helixes support the unwinding activity of Mss116, while the B form DNA does not support its unwinding activity^34^. Thus, the Mss116 protein during its catalytic cycle can discriminate between the A form and B form dsDNA.

The single-stranded poly(A) and poly(C) RNA molecules form ordered coil and helical structures in solution^55^. In solution, the poly(U) RNA samples random coil conformations^55^. On the other hand, the structure of single-stranded poly(G) in solution remains unknown^55^. The inability of poly(G) to support the ATPase activity of DDX1 could be a consequence of its conformation in solution, which is different from a G-quadruplex, and could be different from poly(A), poly(C) and poly(U) homopolymers^55^.

### A blunt-ended dsRNA, dsRNA/DNA, and an ssDNA molecule support the DDX1 protein’s ATPase activity

Single-molecule fluorescence experiments revealed that the Laf-1 DEAD-box protein from *Caenorhabditis elegans* (*C. elegans*) does not bind to blunt-ended dsRNA^56^. In addition, the human ortholog of Laf-1 - the DDX3X protein - which lacked 80 residues from its C-terminal domain was unable to bind to blunt-ended dsRNA molecules; thus, these helixes did not support the C-terminal truncated DDX1 protein’s ATP hydrolysis activity^37^. Moreover, the ATPase activity of a few DEAD-box proteins is stimulated by DNA molecules^26, 27^ ^28^ ^29^ ^30^ ^31^. Here, employing TLC, we investigate whether an ssDNA molecule, a blunt-ended dsDNA/RNA hybrid molecule, or a blunt-ended dsRNA supports the ATP hydrolysis activity of the DDX1 protein (Figure 4, Table1, Table 2).

**Figure 4.**
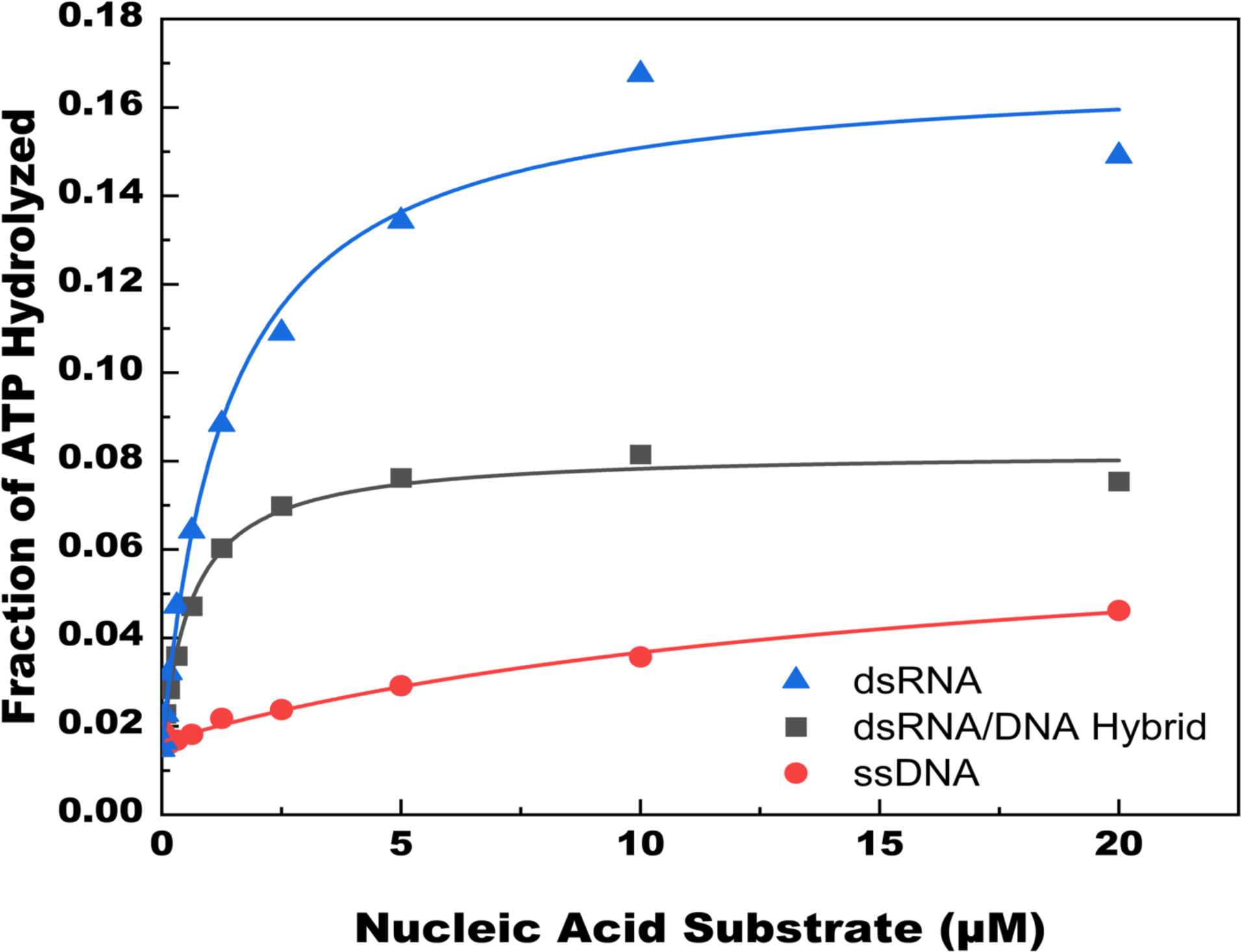
dsRNA, dsRNA/DNA hybrid and ssDNA stimulate the ATPase activity of the DDX1 protein. TLC was used to measure the extent of the DDX1 protein’s ATP hydrolysis in the presence of a blunt ended, dsRNA, blunt-ended, dsRNA/DNA hybrid, or an ssDNA (Table 1). The graphs represent the dependence of the DDX1 protein ATPase activity on the dsRNA, dsRNA/DNA and ssDNA concentrations. Equation 1 was used to fit the data and determine the K_m_ and the maximum fraction of ATP hydrolyzed. The average values for K_m_ and the maximum fraction of ATP hydrolyzed obtained from at least two independent experiments are shown in Table 2. The dependence of DDX1 ATP hydrolysis versus dsRNA concentration is shown in blue triangles, dsRNA/DNA hybrid in black squares, and ssDNA in red circles.

The dependencies of the DDX1 protein’s ATP hydrolysis activity in presence of various concentrations of nucleic acid molecules show that the blunt-ended dsRNA, a blunt ended dsDNA/RNA hybrid, and an ssDNA molecule stimulate the ATP hydrolysis activity of DDX1. However, the extent of the stimulation of the DDX1 protein’s ATP hydrolysis activity is significantly reduced in the presence of an ssDNA molecule and partially reduced in the presence of a dsRNA/DNA hybrid molecule (Figure 4, Table 2).

## CONCLUSIONS

Our experimental results show that the DDX1 protein hydrolyzes ATP and deoxy-ATP only in the presence of RNA and is unable to hydrolyze other nucleosides. The ATPase activity of the DDX1 protein is stimulated by ssRNA molecules, a blunt-ended dsRNA, blunt-ended hybrid dsRNA/DNA, and ssDNA molecule. However, the ssDNA molecule stimulates the ATPase activity of the DDX1 protein significantly less than all the interrogated RNA molecules except for a 10-nucleotide long poly(G) molecule. The poly(G) molecule interrogated here, which is unable to form a G-quadruplex, does not support the ATPase activity of DDX1.

## ASSOCIATED CONTENT

### Accession Code

Human DDX1: Uniprot ID Q92499, human DDX3X: Uniprot ID O00571, human DDX43: Uniprot ID Q9NXZ2*, S. cerevisiae* Dbp5p: Uniprot P20449, human Dbp5p (DDX19): Uniprot ID Q9NUU7, *S. cerevisiae* Dbp9p: Uniprot Q06218, *Pisum sativum* p68: Uniprot ID Q9LKL6, *S.cerevisiae* Mss166: Uniprot ID P15424, *C. elegans* Laf-1: Uniprot ID D0PV95, *S.cerevisiae* Dhh1: Uniprot ID P39517, *T. thermophilus* Hera: Uniprot ID O07897.

---

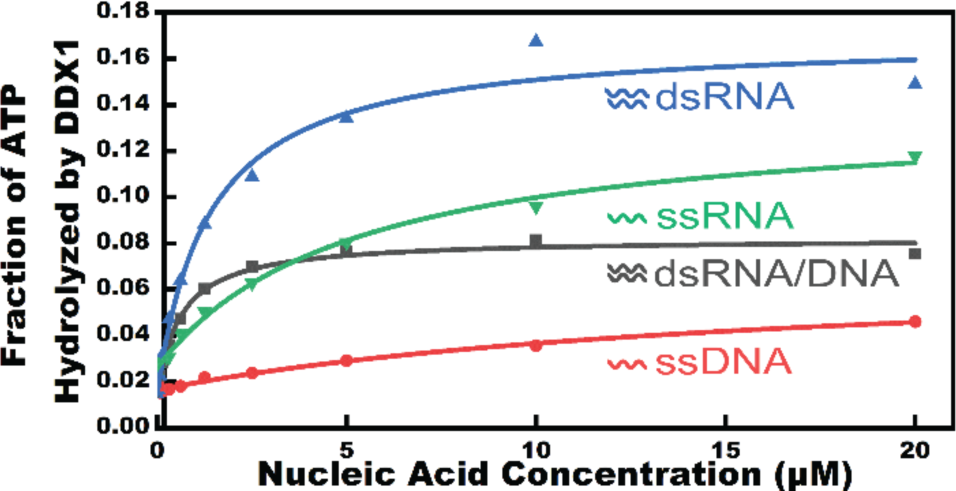
FOR TABLE OF CONTENTS USE ONLY.

